# A Putative Single-Locus Determinant of the Suppressed *In Ovo* Virus Infection (SOV) Trait in *Apis mellifera*

**DOI:** 10.64898/2026.05.28.728461

**Authors:** Regis Lefebre, Bart J. G. Broeckx, Lina De Smet, Merel Braeckman, Aleš Gregorc, Luc Peelman, Dirk C. de Graaf

## Abstract

Today, the deformed wing virus (DWV) can be considered as one of the major causes of global elevated western honey bee colony losses (*Apis mellifera*). Virus transmission may occur horizontally between individuals of the same generation, but also vertically from parents to offspring. The recently defined heritable ‘suppressed *in ovo* virus infection’ (SOV) trait describes the absence of viruses in pooled drone eggs of a queen, associated with significant lower DWV prevalence and viral loads in the subsequent developmental offspring stages. By definition, the trait reflects the absence of vertical virus transmission from SOV-positive (SOV^+^) queens themselves to their offspring. However, the genetic basis influencing this heritable virus resilience has not been identified yet. In this study, we aimed to identify SOV-associated genetic marker(s) or loci in the honey bee genome through genome-wide variant comparison of 44 DWV-positive and 44 DWV-negative drone pupae descendent from an artificially created hybrid SOV^+^/SOV^-^ colony. After whole genome sequencing (WGS), variant calling, and genotype-phenotype association analysis by means of single marker tests and elastic net regression, one variant in a locus of 241.246 bp on chromosome 7 that contained 17 other highly SOV-associated variants classified 68,2% of the drone phenotypes correctly. These results may support the potential application of marker-assisted selection (MAS) strategies targeting reduced vertical virus transmission in honey bees.

## 1. Introduction

Deformed wing virus (DWV) is one of many viruses infecting honey bees, but is probably the most extensively studied one due to its association with its vector, the parasitic mite *Varroa destructor,* and its strong link to honey bee colony losses (1–7). Up to now, three main genotypes of DWV have been described: DWV-A, DWV-B, and DWV-C (4, 8). Over the past decades, epidemiological studies have observed a decrease in the prevalence of DWV-A in both *Varroa*-treated and untreated colonies, while DWV-B has become the dominant strain (8, 9). The emergence of DWV-B has been attributed to its higher transmission efficiency, higher virulence and faster replication in adult hosts, and overall global beekeeping and bee migration practices (9–12). Despite being identified as phylogenetically distinct from DWV-A and DWV-B within the DWV species complex, DWV-C has been detected only sporadically and at low levels in a limited number of surveys (9, 13). Altogether, DWV-B, including any recombinants with DWV-A, may be considered as one of the major causes of elevated overwinter honey bee colony losses (10, 14).

Transfer of viruses between individual bees of the same generation is known as horizontal transmission, which may happen through either direct or indirect contact (6, 15–17). Examples of such transmission routes for honey bee viruses are fecal-oral transmission, bodily contact, handling of contaminated objects, trophallaxis, food contamination, brood cell cleaning, and transmission via the *Varroa* mite (15, 17). For a long time, it was unclear whether the *Varroa* mite purely acted as a vector facilitating DWV transmission, or whether DWV could infect and actively replicate in the mite (4, 5, 18). Recent findings showed that, next to other important honey bee viruses, DWV-B is able to infect and replicate in *Varroa*, and that the mite thus acts as a biological vector (4, 5). Whether or not this holds equally true for DWV-A remains unsure (2, 4, 5).

Vertical or transgenerational transmission of honey bee viruses by queens and drones to offspring has been demonstrated to occur via eggs and semen, respectively (16, 17, 19–23). Ravoet et al. (2015) reported a sanitary screening method to investigate the portion of vertical virus transmission from honey bee queens and drones to their offspring in an attempt to increase the overall success rate of queen rearing in the Belgian queen breeding program (21). Therefore, queen breeders were asked to pool 10 worker eggs per test queen, which were screened by RT-PCR assays for the presence of multiple honey bee viruses, including acute bee paralysis virus (ABPV), aphid lethal paralysis virus (ALPV), black queen cell virus (BQCV), DWV, lake Sinai virus (LSV), sacbrood virus (SBV) and *Varroa destructor* Macula-like virus (VdMLV) (21). For the first time, this study reported the vertical transmission of ALPV, LSV, and VdMLV in honey bees, and promoted the implementation of similar sanitary screenings in breeding programs to increase the overall queen rearing success (21).

Further studies showed that colonies headed by virus-free egg-laying queens exhibit fewer and less severe DWV infections across nearly all developmental stages of both drone - and worker offspring (24, 25). Hence, de Graaf et al. (2020) estimated the heritability of this trait (h^2^ = 0,25) and further confirmed that colonies headed by so-called ‘suppressed *in ovo* virus infection (SOV)’ trait -positive queens (SOV^+^) showed an increased resilience to DWV-A and DWV-B in almost all developmental offspring stages, with a particularly pronounced effect in drones (20, 24, 25). Thus, the viral status of unfertilized drone eggs is a good indicator of the portion of vertical virus transmission of the queen herself to the offspring (20, 24–26). Concisely, the current definition of the SOV trait comprises the absence of viral infections in sampled drone eggs, associated with significant lower DWV prevalence and viral loads in subsequent developmental stages (24), thereby reflecting the absence of vertical virus transmission from queens themselves to their offspring (20, 24, 25). Due to its moderate heritability and its beneficial effects in terms of colony health, implementation of the SOV-trait in breeding programs was highly recommended (24, 25).

Antiviral defense in honey bees is mediated through the RNA interference (RNAi) pathway, signal transduction cascades triggered by pathogen-associated molecular patterns (PAMPs) such as the Janus kinase and Signal Transducer and Activator of Transcription (JAK-STAT), immune deficiency (Imd) and Toll pathways, generation of reactive oxygen and nitrogen species (ROS/RNS), and restriction of viral proliferation through apoptosis (27–33). As the largest portion of honey bee viruses are positive-strand RNA viruses (+ssRNA) requiring dsRNA intermediates during replication, the RNAi pathway forms the most important antiviral response in honey bees (6, 27). Apart from these antiviral defense mechanisms, any additional heritable basis affecting the outcomes of viral infections in honey bees, and their resilience towards viruses, has not been identified yet (27, 34). However, quantitative trait loci (QTL) mapping studies have already been performed for Israeli acute paralysis virus (IAPV) susceptibility, and SBV resistance (34, 35).

In this study, we aimed to identify SOV-associated genetic markers or loci in the honey bee genome through haploid genome-wide variant comparison of DWV-B positive and DWV-B negative drones descending from an artificially created hybrid SOV^+^/SOV^-^ colony. Creation of a hybrid colony allows to bypass false positive associations due to population stratification, and provides phenotypic groups of drones that are genetically related, except for the segregated phenotype-associated variants (36–39). After genome sequencing of the drones from both phenotypic groups, variants were called and individually analyzed for their association with the samples’ DWV-B viral infection status. Since traditional single-marker tests often lack the capacity of testing multiple variants collectively for their association with the phenotype of interest, we subsequently applied elastic net regression for joint variant modelling (40–42). Finally, the predictive performance and performance parameters of the rendered model(s) were evaluated.

## 2. Materials and methods

### 2.1. Honey bee colony selection and creation of hybrid SOV^+^/SOV^-^ colonies

Virgin virus resistant (SOV^+^) *Apis mellifera carnica* queens, reared from four different SOV^+^ pedigrees through standard queen rearing (43), were each inseminated with the semen of a single drone from a virus sensitive (SOV^-^) ssp. *carnica* line (P-crossing) (38). Single drone inseminations (SDI) were performed in line with the standard methods for instrumental insemination of *Apis mellifera* queens (44). For each single drone, 1,5 µL semen was diluted with 0,5 µL SDI-buffer (0,2 M NaCl; 5 mM glucose monohydrate; 0,67 mM L-lysine; 0,57 mM L-arginine; 0,68 mM L-glutamic acid; 0,02 M Trizma HCl; 0,03 M Trizma base and 2,5 mg/mL dihydrostreptomycin) and inseminated in the oviduct of the CO_2_- anaesthetized virgin queen. After introduction of these single drone-inseminated queens in small Ein Waben Kästchen (EWK) nucleus hives, new hybrid SOV^+^/SOV^-^ queens were produced through standard methods of queen rearing (F1) (38, 43). Again, these virgin queens were introduced in small EWK nucleus hives, but were allowed to mate naturally. After winter, surviving hybrid SOV^+^/SOV^-^ queens were transferred to standard Simplex hives. Finally, drone brood (F2) being produced by hybrid queens during the bee season has been sampled by introduction of drone cell comb frames.

### 2.2. Prescreening of hybrid colonies

Seventeen days after introduction of the drone cell comb frames into the colonies with a hybrid SOV^+^/SOV^-^ queen (F1), the capped drone brood was collected and frozen at -80°C. From each frame, ten randomly selected non-*Varroa*-infested drone pupae (red eyes, 17 days old) were sampled and prescreened for DWV by Reverse Transcription Quantitative Polymerase Chain Reaction (RT-qPCR).

Therefore, total RNA was extracted from individual whole drone bodies through PowerLyzer® 24 homogenization in the presence of 1 mL QIAzol^TM^ Lysis Reagent (QIAGEN), zirconia beads and 3 metal beads at 30 Hz for 1 minute. After addition of 200 μL chloroform and incubation at room temperature (RT) for 5 minutes, the samples were centrifuged at 12.000g, and the upper phases used for RNA extraction using the RNeasy® Lipid Tissue Mini Kit (QIAGEN), following the manufacturer’s instructions. The RNA was finally eluted in 50 μL DNase/RNase free water. Complement DNA (cDNA) synthesis was performed with the RevertAid H Minus First Strand cDNA Synthesis Kit (Thermo Scientific), using random hexamer primers and 5 µL RNA as template. Next, the cDNA was diluted 5x and stored at -20°C. Quantitative PCR for relative quantification of the DWV load was performed using the Platinum^TM^ SYBR^TM^ Green qPCR SuperMix-UDG (Thermo Scientific) using a deformed wing virus FAM (DWV-FAM) primer set detecting the DWV complex (Supplementary Table S1). Cycling settings were 2 minutes at 50°C; 2 minutes at 95°C; and 40 x (15 sec. 95°C; 20 sec. 58°C; 30 sec. 72°C). No template controls (NTCs) and melt-curve dissociation analyses (65°C – 95°C) were included to check for target specificity.

### 2.3. Drone sampling, DNA/RNA extractions, and determination of relative DWV-B load by RT-qPCR

Based on the prescreening results, a total of 291 drone pupae with red eyes (17d) were sampled from the preserved drone brood of a single selected colony and screened for DWV-B. From all 291 collected pupae, genomic DNA and RNA were simultaneously extracted by using the QIAGEN AllPrep DNA/RNA/Protein Kit (QIAGEN). More specifically, individual pupal bodies were homogenized with a PowerLyzer® 24 in the presence of 987 µL RLT Plus buffer, 3 metal beads, sterile zirconium beads and 13 µL β-mercapto-ethanol. One third of the obtained whole body lysate was then used for subsequent RNA and DNA extraction by using AllPrep DNA spin columns, according to the manufacturer’s protocols. Per sample, RNA was eluted twice from the RNeasy spin columns (2x 50 µL) with DNase/RNase free water. DNA was eluted from the All prep DNA spin columns with 80 µL EB buffer.

Subsequent cDNA synthesis and relative quantification of the DWV-B load by qPCR were performed as described in 2.2., but using a DWV-B primer set (Supplementary Table S1). Absolute quantification was not performed, as subsequent creation of DWV-B^+^ and DWV-B^-^ groups could be accomplished based on Ct values only. Forty-four drone pupae showed Ct values lower than 31 on qPCR and were considered at least moderately infected by DWV-B (> 10^5^ viral particles/drone) (20). To obtain equal groups for comparative variant calling, 44 samples were randomly chosen from the 59 DWV-B negatives (by a random number picker). For all 88 selected samples, separate qPCRs with β-actin, ABPV, BQV, and SBV as targets were performed as described above in 2.2., to assure presence of cDNA and absence of other viruses, respectively (Supplementary Table S1).

### 2.4. Whole genome sequencing (WGS) and variant calling

#### 2.4.1. DNA extraction and sample preparation

From all eighty-eight selected drones (44 DWV-B negative; 44 DWV-B positive), genomic DNA was obtained from the whole body as described in 2.3. For purification, the EB buffer-solved DNA was precipitated with 5 M ammonium acetate and 100% ethanol, washed with 70% ethanol and resuspended in 20 µL DNase/RNase free water. Concentrations and optical densities (ODs) were measured by using Nanodrop spectrophotometry.

Genomic DNA samples were sent to BGI Genomics for whole genome sequencing (WGS). For this, at least 400 ng of DNA at a minimal concentration of 8 ng/µL was prepared for each sample and shipped to BGI Genomics on dry ice. After sample quality checks (QC) by BGI, normal DNBseq DNA libraries were constructed, and library QC were performed with electropherograms.

#### 2.4.2. Whole genome sequencing (WGS), sequencing data analysis and mapping

The used sequencing platform by BGI was BGISEQ-500 WGS, implementing DNA Nano Ball sequencing (DNB Seq) with paired end read lengths of 150 bp. The SOAPnuke (Version: 2.2.1) tool was used for adapter trimming, low quality reads trimming and contiguous N bases trimming. The filtered data, called ‘clean data’, was used for subsequent analysis.

For each drone, clean reads were mapped to the reference genome GCF_003254395.2_Amel_HAv3.1 by the short reads mapping software BWA (Burrows-Wheeler Aligner) (Version: 0.7.17) in “mem” module. Samtools (Version: 1.9) was used to shift the sequence alignment map (SAM) result to binary alignment map (BAM) format and to fixmate, markdup and sort the BAM files. Samtools and Qualimap2 (Version: 2.2.1) were used to calculate coverages, sequencing depths and plotting of the alignments (45). After this, the processed BAM files were ready for variant detection.

#### 2.4.3. Variant calling and filtering

The Genome Analysis Toolkit (GATK) was used to call SNPs and indels. More specifically, GATK4 (Version: 4.2.6.1) was used to detect and call SNPs and indels for each sample. As drones were sequenced, “ploidy” was specified as being 1 in GATK HaplotypeCaller. Low quality SNPs were removed if they met one of the following conditions (GATK VariantFiltration): QD < 2,0 || FS > 60,0 || MQ <40,0 || MQRankSum < -12,5 || ReadPosRankSum < -8,0. Low quality indels were removed if they met one of the following conditions (GATK VariantFiltration): QD < 2,0 || FS > 200,0 || SOR > 10,0 || MQRankSum < -12,5 || ReadPosRankSum < -8,0.

### 2.5. Single marker analyses and elastic net regression

Final data was stored as standard Variant Call Format (VCF) files, in which called variants (SNPs and indels) were grouped per chromosome. R (version 2024.04.1+748) and RStudio were used for single marker analyses (SMA) and penalized elastic net regression. After different filtering steps (Supplementary Table S2), the remaining SNPs and indels were tested by Fisher’s Exact Tests with Bonferroni correction.

For elastic net regression, unaltered variables were all SNPs and indels with a call for all 88 samples (N = 1.149.592) and the phenotype of the samples (DWV-B negative (0); DWV-B positive (1)). For optimal lambda and alpha selection, alpha was altered from 1 to 0,1 in steps of 0,1 and lambda was tested repetitively for 100 different values. For each value of alpha, the number of included variants and percentages of correct classification by the models at λ_min_ and λ_1.se_ were determined.

## 3. Results

### 3.1. Creation of hybrid SOV^+^/SOV^-^ colonies and subsequent screening for DWV(-B)

A total of nine single drone-inseminated ‘homozygous’ SOV^+^ queens produced worker brood after SDI (at least one per SOV^+^ pedigree). From these nucleus colonies, twelve new ‘hybrid’ SOV^+^/SOV^-^ queens were produced (F1). After winter, six of these hybrid queens survived, of which five (from three different lines) produced drone brood (F2) during the bee season, denoted as colonies A-E (Figure 1).

**Figure 1.**
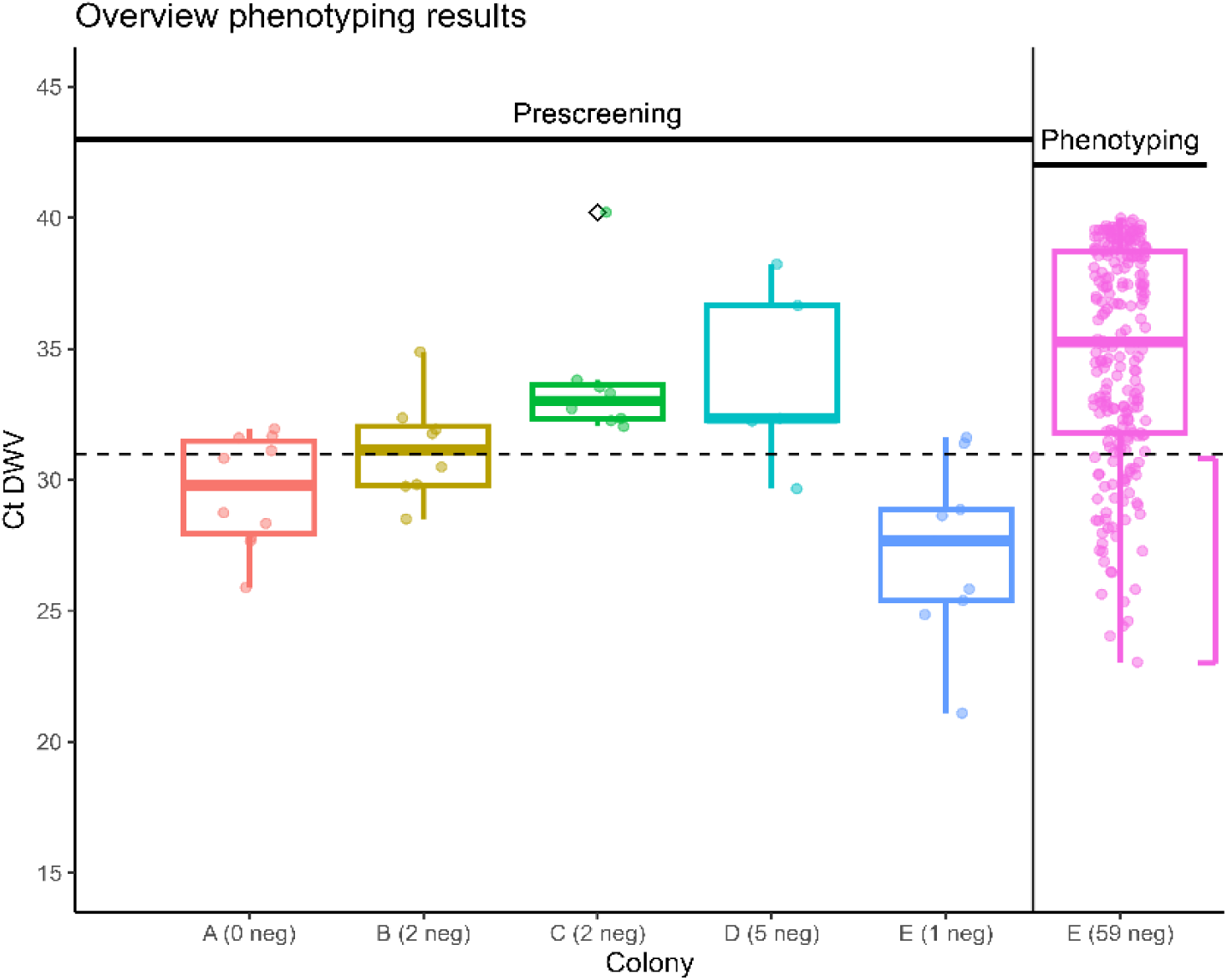
Prescreening of the 5 hybrid colonies (A-E), and subsequent phenotyping of 291 pupae of colony. **E.** For each colony, the number of negative samples in the respective (pre)screening is given (‘neg’). The dashed line represents Ct = 31. The bracket indicates the 44 selected DWV-B -positive drones (Ct < 31) among the phenotyped pupae of colony E. Lower Ct values correspond to higher viral loads. Samples were set negative if the Ct was not reached at the final cycle (i.e. 40 here).

Prescreening these five hybrid queens enabled us to select colony E, which showed the highest segregation in relative DWV load in the offspring drone pupae (Figure 1). In that colony, 10% of the prescreened drone pupae were completely DWV-negative (1 out of 10), while the DWV-positive samples showed the highest average loads (avg. Ct = 27,3) when compared to the other colonies’ drones. Based on this prescreening, colony E was selected for subsequent screening of F2 drones for their relative DWV-B load. Fifty-nine of the 291 screened drones (20,3 %) were DWV-B negative (no Ct in duplicates) (Figure 1). Forty-four drone pupae showed Ct values lower than 31 on qPCR and were considered at least moderately infected by DWV-B (> 10^5^ viral particles/drone) (Figure 1). Subsequent qPCRs targeting ABPV, BQV, and SBV revealed absence of these viruses in the final 88 selected samples.

### 3.2. Whole genome sequencing and mapping

After sequencing, a total of 4.765.417.618 and 4.672.791.826 (98%) raw/clean reads were obtained, resulting in an average of 54.152.473 and 53.099.907 raw/clean reads per sample, respectively (Supplementary Figure S1a, Supplementary Table S3). Moreover, an average of 52.846.249 clean reads per sample were successfully mapped (mapping rate: 99,5%; Figure S1a, Table S3). An overall average sequencing depth of 34,7x was obtained (Figure S1b, Table S3). Among all samples, at least 97,2% and 94,6% of the target nucleotides were covered ≥5x and ≥10x, respectively (averages: 97,6% and 96,4%; respectively) (Figure S1c and S1d, Table S3).

### 3.3. Variant calling and single marker analyses (SMA)

After variant calling, a total of 2.203.547 different SNPs and indels were found among all 88 sequenced samples. SMA were performed to determine individual associations of SNPs and indels with the phenotype of interest (DWV-B positive vs. DWV-B negative). After different filtering steps (Supplementary Table S2), a total of 1.221.554 remaining SNPs and indels were tested for their association with DWV-B by Fisher’s Exact Tests. A Manhattan plot was constructed for all filtered SNPs and indels, and variants were grouped by chromosome (Figure 2). None of the 1.221.554 SNPs and indels reached the Bonferroni-corrected genome-wide significance threshold (α/number of tested SNPs). However, as a reference, the most significant SNP and indel per chromosome are depicted in Supplementary Table S4.

**Figure 2.**
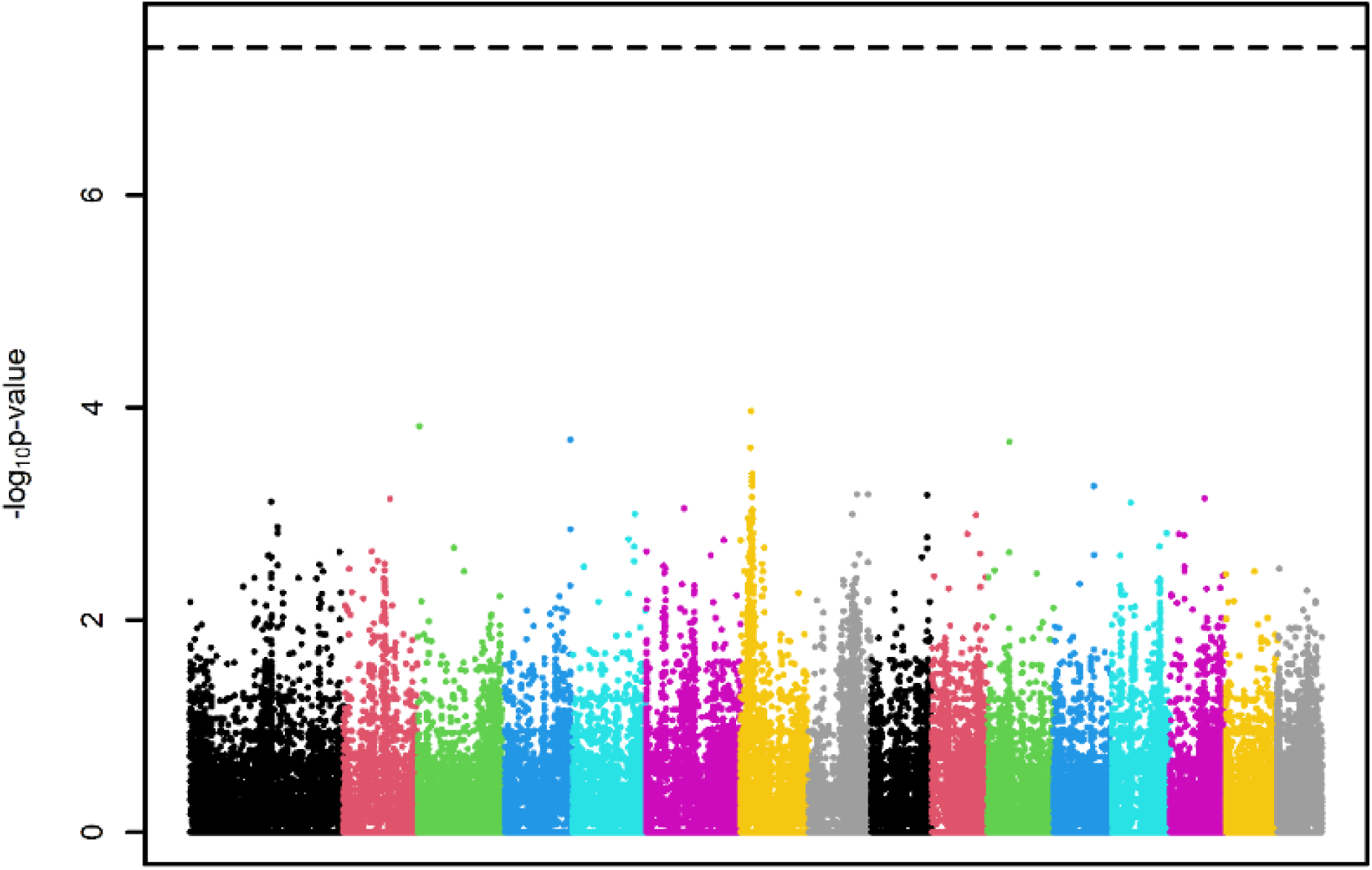
SMA Manhattan plot of uncorrected p-values of all tested SNPs and indels (N = 1.221.554). P-values were obtained using Fisher’s Exact Tests. The dashed line depicts the threshold for significance (Bonferroni-corrected α = 0,05/ 1.221.554 variants). SNPs and indels are grouped by chromosome (= Linkage Group; colors) and ordered by position. From left to right: chromosomes 1 to 16.

Of all 30 single marker tests with uncorrected p-values ≤0,001, 7 indels and 11 SNPs (18/30 = 60%) are located on a 241.246 nucleotide-wide locus on chromosome 7 (NC_037644.1) (Table 1, Figure 3, Supplementary Figure S2 and Table S5). Genes occurring in this locus are GAS2-like protein pickled eggs (LOC411586), take-out-like carrier protein (JHBP-1), zinc finger protein 470 (LOC725174), enolase-phosphatase E1 (LOC100577098), protein kinase C-binding protein NELL1 (LOC411748), alpha-N-acetylglucosaminidase (LOC551437), mitochondrial amidoxime-reducing component 1 (LOC100577026), 4-nitrophenylphosphatase (LOC551405), and multiple uncharacterized genes (LOC100578137, LOC100577132, LOC100577221, LOC413153, LOC113218887, LOC102654639) (Figure 3, Supplementary Table S5).

**Figure 3.**
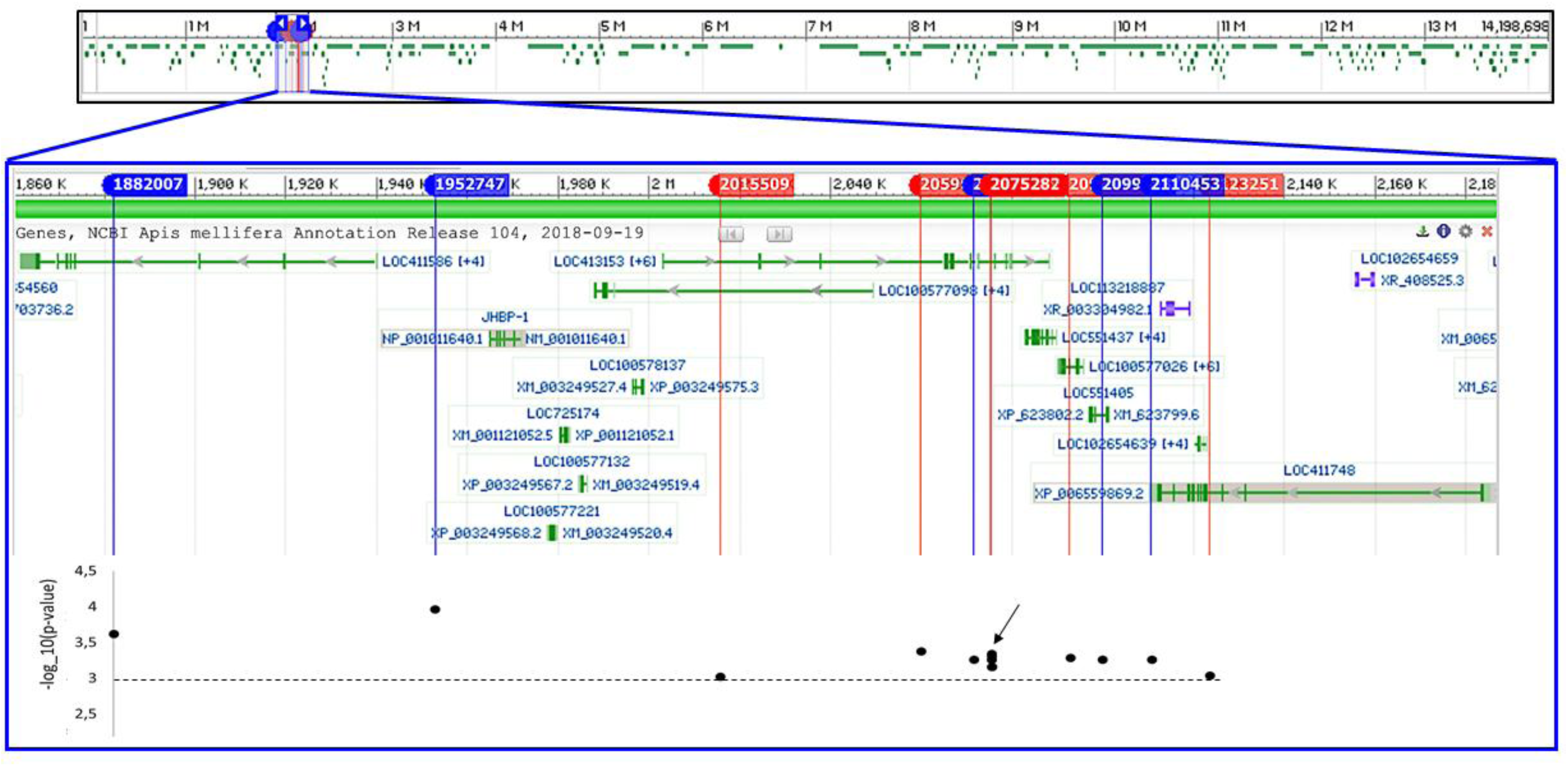
Overview of the 18 variants with uncorrected p-value ≤0,001 in the 241.246 nucleotide-wide locus on chromosome 7. Starting- and ending position of the range on reference genome Amel_HAv3.1 is 1.882.005 and 2.123.251, respectively. Indels are depicted in blue, SNPs in red. Below, the -log_10_ of the *p*-value of the SMA for each of the variants is shown, with the dashed line indicating the threshold of *p* = 0,001 (-log_10_(0,001) = 3). Of the 18 significant variants, seven SNPs and one indel are all located in the same 8th intron of the uncharacterized LOC413153 gene (arrow). Figure made with NCBI’s Genome Data Viewer.

**Table 1.** Overview of all variants with uncorrected p-value ≤0,001 for the single marker analyses (Fisher’s Exact Tests). Positions and ranges correspond to the reference genome Amel_HAv3.1. In total, 30 variants (17 indels and 13 SNPs) showed an uncorrected p-value ≤0,001 for the single marker analyses. Eighteen out of 30 (60%) variants were located on a 241.246 nucleotide-wide locus on chromosome 7 (asterisk).

| Chromosome | Indels / SNPs | Position(s) or range | Total variants |
| --- | --- | --- | --- |
| 1 | 1/0 | 13.917.351 | 1 |
| 2 | 1/0 | 10.883.424 | 1 |
| 3 | 1/0 | 922.528 | 1 |
| 4 | 1/0 | 12.629.047 | 1 |
| 6 | 1/0 | 6.778.691 | 1 |
| 7 | 7/11 | 1.882.005 – 2.123.251 | 18* |
| 8 | 1/1 | 9.616.930/ 11.459.866 | 2 |
| 9 | 1/0 | 11.404.115 | 1 |
| 11 | 1/0 | 6.706.915 | 1 |
| 12 | 1/0 | 8.161.396 | 1 |
| 13 | 1/0 | 5.349.407 | 1 |
| 14 | 0/1 | 6.252.297 | 1 |
| **Total** | **17/13** |  | **30** |

Of the 18 variants located in the abovementioned locus, nine SNPs and two indels were found to be located in the introns of an 85.477 nucleotide-long uncharacterized gene (LOC413153) (Figure 3, Supplementary Table S5). Moreover, seven SNPs and one indel were all located in the same 8^th^ intron of this gene (arrow in Figure 3).

### 3.4. Elastic net regression

During elastic net regression model optimization by leave-one-out cross-validation, the unaltered variables in this study were all 1.149.592 SNPs and indels with a call for all 88 samples (that is 94,1% of the 1.221.554 filtered variants in the SMA), and the phenotype of the samples (DWV-B^-^ (0); DWV-B^+^ (1)). For optimal lambda and alpha selection, alpha was altered from 1 to 0,1 in steps of 0,1 and lambda was tested repetitively for 100 different values (Figure 4, Supplementary Table S6).

**Figure 4.**
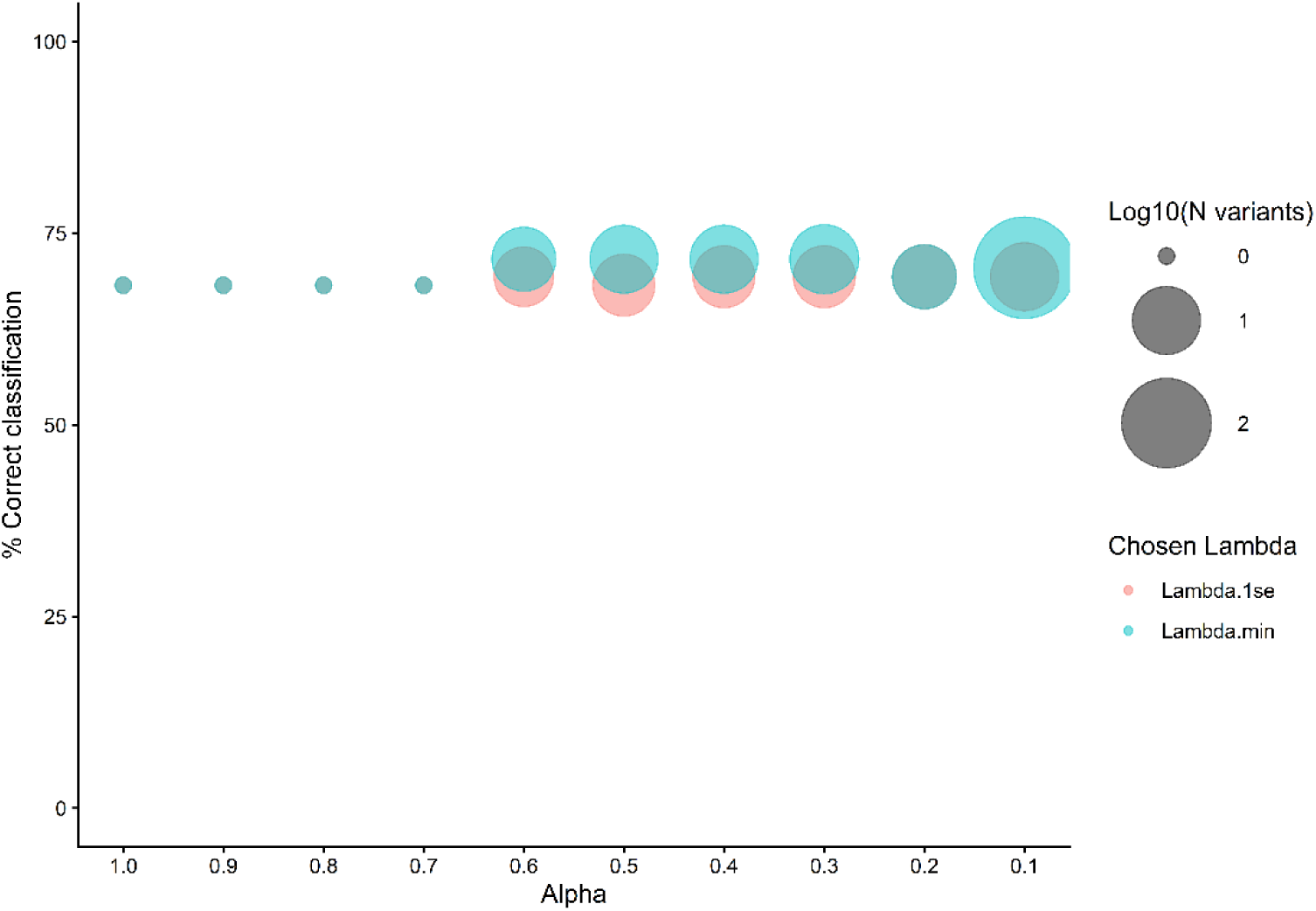
Overview of elastic net regression analyses. In these analyses, all 1.149.592 SNPs and indels with a call for all 88 samples were considered as the unaltered predictor variables, and the phenotype of the samples (DWV-B^-^ (0); DWV-B^+^ (1)) as binary outcome variable. For optimal lambda and alpha selection, alpha was altered from 1 to 0,1 (in steps of 0,1), and lambda was tested repetitively for 100 different values. For each value of alpha, the number of included variants and percentage of correct phenotype classification by the models at λ_min_ (blue) and λ_1.se_ (pink) are represented.

Although some models with lower alpha values rendered higher percentages of correct phenotype prediction by integrating more genetic variants as predictors, the differences in predictive accuracy compared to models with only one variant (α= 1 to 0,7) was rather minimal (Figure 4). For example, the model at α= 0,6 and λ_min_ = 0,292 comprised seven genetic variants, but predicted only 3,4% of the phenotypes more correctly compared to models with only one genetic variant (that is 71,6% at α= 0,6 and λ_min_ vs 68,2% at α= 1 to 0,7, respectively) (Figure 4). Therefore, the model with α= 1 (LASSO only) at λ_min_ = λ_1se_ = 0,192 with only one single genetic variant as predictor was retained for further analysis (Figure 4). In this model, the intercept and estimate of the single retained variant equaled -0,0715 and 0,0797, respectively, identifying it as being a risk variant (estimate > 0). The specificity, sensitivity, negative predictive value (NPV) and positive predictive value (PPV) of this single-variant model all equaled 68,2% (Table 2). The single significant predictive variant in the abovementioned model is an intergenic indel at position 1.952.747 on Linkage Group 7 (LG7; NC_037644.1; Amel_HAv3.1). This was also the most significant variant within the 241.246 nucleotide-wide locus on chromosome 7 in the SMA (Figure 3, Supplementary Table S5).

**Table 2.** Classification table comparing the predicted phenotypes (‘predicted by model’) relative to the observed phenotypes (‘phenotypic group (truth)’) of the 88 drones. 68,2% of the samples were correctly classified (60/88).

| Predicted by model | Phenotypic group (truth) |  |  |
| --- | --- | --- | --- |
|  | 1 (DWV-B<sup>+</sup>) | 0 (DWV-B<sup>-</sup>) |  |
| 1 (DWV-B<sup>+</sup>) | 30 | 14 | NPV: 68,2% |
| 0 (DWV-B<sup>-</sup>) | 14 | 30 | PPV: 68,2% |
| Total | 44 | 44 |  |

In relation to the remaining 17 SMA-significant variants located along this 241.246 nucleotide-wide locus, Supplementary Figure S2 and Table S7 represent the alleles associated with the variant type alleles Vt_1_ and Vt_2_ of the above-described indel (NC_037644.1_1952747). To this purpose, multiple variant type alleles were considered as one Vt allele group.

## 4. Discussion

Together with and mediated by *V. destructor*, viral infections contribute significantly to the globally increasing honey bee colony mortality rates (6, 20, 46). Although honey bee viruses commonly occur in colonies without pathogenic symptoms, *Varroa* mite infestations may significantly change the prevalence, virulence and distribution of these viruses in the considered honey bee population (46). Contrary to *Varroa* resistance, any forms of resilience to viral infections are rarely implemented in honey bee breeding programs.

However, one way of screening honey bee colonies for vertical virus transmission is by scoring them for the ‘suppressed *in ovo* virus infection (SOV)’ trait, describing the absence of viral infections in sampled drone eggs, correlated with increased resilience of the colony to viral infections as a whole (24, 47). Therefore, the SOV trait also reflects the queen’s viral resilience and absence of vertical virus transmission to her offspring (20, 24, 25, 47). Although the trait is originally described with pooled drone eggs as target samples (20, 24), it has been shown that colonies expressing the SOV trait display fewer and less severe DWV infections in most developmental offspring stages, especially in drones. More specifically, de Graaf et al. (2020) studied the proliferation of the SOV trait on the DWV infection levels in drone eggs, larvae, pupae and adults sampled from 4 SOV^+^ colonies versus 4 SOV^-^ colonies, and showed a strong beneficial effect of the SOV trait on the male bee caste in terms of viral prevalence and load, especially in larval and pupal stages (24). Regarding our search for genetic markers in the honey bee genome associated with the SOV trait, screening individual drone eggs was unfeasible due to the low amounts of extractable DNA and RNA. Alternatively, we sampled and phenotyped individual drone pupae, postulating that the molecular basis and expression of the SOV trait is equally well reflected in this developmental caste and stage, whereas the haploid genomes significantly ease further genomic analyses.

Disregarding vertical transmission from queens and drones to their offspring, transmission of honey bee viruses to the developing brood may also occur horizontally through contamination of cells cleaned by infected workers, infestation by *Varroa* mites, and consumption of contaminated food (16, 17). By consequence, any differences in these horizontal transmission routes may have influenced the viral loads in the phenotyped offspring drones of the selected hybrid colony. We do admit that *in vitro* rearing of sampled eggs in a sterile laboratory environment would have provided more control on these horizontal transmission routes. However, several measures of phenotype constriction were considered to reduce the influence of confounding factors. First, although the prescreening rendered different DWV infection patterns among the five artificially created hybrid SOV^+^/SOV^-^ colonies, it should be noted that the final comparative GWA analysis would and did only occur between pupal samples of the same hybrid colony. Moreover, all DWV(-B) screened drone pupae were sampled from the same side of a single capped drone brood frame of the selected hybrid colony. As the *Varroa* mite is the main biological vector for DWV (4), determination of the DWV(-B) viral status was only performed for drone pupae uninfected by *Varroa*. In addition, as mixed viral infection patterns may bias the GWAS analysis, the sampled drone pupae of the selected hybrid colony were screened for three other viruses than DWV-B, namely ABPV, BQCV and SBV. Although we realize that these four viruses represent only a fraction of the entire honey bee virome, they have been reported to be the most prevalent ones in Belgium according to a population-wide screening of more than 500 colonies from 155 Belgian beekeepers (48).

Although one would expect two clearly segregated phenotypic groups after determination of the relative DWV-B viral load in the sampled drone pupae from the hybrid SOV^+^/SOV^-^ colony, it should be highlighted that the SOV trait is associated with reduced virus prevalence and viral loads in the colony, and not with total absence of it. Stated differently, colonies headed by SOV^+^ queens may still have DWV titers in part of the larval, pupal or adult offspring population, but in general, the prevalence and loads will be lower. We hypothesize that the observed distribution of relative DWV-B load in the 291 screened drone pupae is the result of the combination of lower, but not null, and higher prevalence and viral loads from the SOV^+^- and SOV^-^ background of the hybrid colony, respectively. Anyhow, by sampling drones from both distributional extremes (moderately-highly positive vs. total absence), we increased our chances of including offspring drones either carrying SOV^-^ or SOV^+^-associated alleles, respectively.

In this study, absolute virus quantification was not performed, as subsequent creation of DWV-B^+^ and DWV-B^-^ pupal groups could be performed based on Ct values only. However, based on our experience with qPCR-based viral quantification in honey bees using calibration curves, we estimate a viral load in the considered sample type between 10^5^ and 10^6^ per drone at Ct = 31. Consequently, all selected DWV-B positive samples contained DWV-B loads higher than 10^5^. In addition, supplementary viral screenings showed that all selected samples, including the 44 DWV-B positive drones, were free of ABPV, BQCV and SBV. These findings are in line with similar screening results in other studies (47, 49), and match the yearly results of viral screenings in the Flemish Bee Breeding Program, wherein ABPV, BQCV and SBV infections in pooled drone egg samples repetitively show low prevalence among tested queens. The hybrid elastic net regression combines the strengths of ridge - and lasso regression, and is used in cases with many variables whereof it is unknown whether they are useful or not (42). In addition, the method deals especially well with situations comprising correlations between parameters. When sample sizes (here: N = 88 drones) are relatively small compared to the number of independent variables or parameters (here: 1.149.592 genetic variants), which is often the case in comparable GWAS analyses, ridge regression can improve predictions made from new data by making predictions less sensitive to the training data set (41). While ridge regression still retains all variables, lasso regression may exclude useless variables from equations (parameter shrinkage) (40). By combining lasso and ridge regression, elastic net regression groups and shrinks parameters associated with correlated variables, and leaves them in the equation or removes them all at once.

In the current study, the single significant variant associated with the SOV trait in the retained elastic net model was an intergenic indel at position 1.952.747 on chromosome 7. Regarding the single marker analyses (SMA), this indel was the most significant of all 18 variants within the 241.246 nucleotide-wide locus on chromosome 7. Hence, both the SMA and elastic net regression analysis highlighted the association of this locus and its variants with the SOV trait.

Although all SOV-associated variants highlighted in this GWA study are located within intronic and intergenic regions in the significant locus on chromosome 7, such variants are increasingly being recognized as functionally relevant in the context of gene regulation (50–53). For instance, genetic variations in noncoding regions may influence gene expression through modification of regulatory elements such as enhancers, silencers, or insulators, thereby affecting transcription of nearby genes in *cis* or more distal targets (*trans*). In addition, intronic variants may alter splicing motifs, leading to different transcript structures or abundances. Moreover, variants in noncoding regions may also affect chromatin accessibility, DNA methylation patterns, or the expression of noncoding RNAs, which may all contribute to regulatory variation. Collectively, these mechanisms explain how GWAS-discovered variants outside coding regions can contribute to phenotypic variation in the trait of interest. Given these regulatory mechanisms, characterization of the genes within the associated locus is critical for identifying potential biological mediators of the observed association. The SOV-associated 241.246 nucleotide-wide locus on chromosome 7 contains 8 characterized and 6 uncharacterized genes, whose main function and potential link to viral infections are described below.

*Drosophila melanogaster’*s Gas2-like protein pickled eggs (Pigs) binds both actin and microtubular cytoskeletons, thereby acting as a cytoskeletal crosslinker (or cytolinker) affecting cytoskeleton organization (54, 55). Many viruses use major cytoskeletal networks such as actin filaments and microtubules as cellular migratory transport systems during the early stages of replication (56–59). Moreover, it has been shown that numerous viruses manipulate their host’s cytoskeletal structures near the plasma membrane to facilitate viral entry through endocytosis (60–63).

Take-out-like carrier proteins with juvenile hormone binding motifs, such as juvenile hormone binding protein 1 (JHBP-1), are members of a protein family regulating embryogenesis and stimulating reproductive maturation in adult honey bees (64–66). For instance, some insect JHBPs have been found to serve as transporters of juvenile hormones (JH) to target tissues, preventing molting and metamorphosis by interfering with the ecdysone-induced changes in gene expression patterns (67–72). Studies associating insect JHBPs with viral infections are rather scarce, although one study in *Aedes aegypti* reported that silencing of RpL23 and RpL27, two ribosomal large subunits, increased resistance to Zika virus (ZIKV; *Flaviviridae*) infection (73). Further analysis showed that expression of both ribosomal subunits is regulated by the JH signaling pathway (73).

Enolase-phosphatase E1 (ENOPH1, also noted as MASA) is an enzyme converting 2,3-diketo-5-methylthio-1-phosphopentane to the intermediate 1,2-dihydroxy-3-keto-5-methylthiopentene anion in the methionine salvage pathway (74–76). This pathway consists of six reactions recycling methionine from 5’-methylthioadenosine (MTA), a byproduct of the polyamine synthesis pathway (76, 77). Higher expression of ENOPH1 has been described during dengue (DENV) and chikungunya virus (CHIKV) infection of *Aedes albopictus* cell culture systems, while ENOPH1-knockout mosquito cell lines showed higher CHIKV titers compared to control lines (78–80). Although ENOPH1 is well known to play an important role in cellular stress response through its function in polyamine production (81, 82), these studies also indicate its importance in reducing viral replication in host cells (79).

Alpha-N-acetylglucosaminidase (NAGLU) is a lysosomal enzyme degrading heparan sulfate, a glycosaminoglycan (GAG), by catalyzing the hydrolysis of terminal non-reducing N-acetyl-D-glucosamine residues in N-acetyl-alpha-D-glucosaminides (83). Any disfunction of this enzyme may lead to an accumulation of GAGs (84). Regarding viral infections, surface-present heparan sulfate proteoglycans (HSPGs) are heavily sulfated (SO_4_^-^), resulting in a global negative charge that can interact with human pathogenic viruses, or may even act as direct viral entry receptors (85–90). For instance, cell surface heparan sulfate (HS) has been characterized as an important Dengue virus receptor in different human model systems (91).

To the best of our knowledge, no studies have yet reported on the role of protein kinase C-binding protein Neural EGFL Like 1 (NELL1), Zinc finger protein 470 (ZNF470), mitochondrial amidoxime-reducing component 1 (MTARC1 or mARC1), and 4-nitrophenylphosphatase (PNPPase), or their link to viral infections in insects.

Further investigation of the 6 uncharacterized genes located in the 241.246 nucleotide-wide locus on chromosome 7 with the Basic Local Alignment Search Tool (BLAST) revealed two non-coding sequences (LOC113218887 and LOC102654639), three uncharacterized protein coding genes not showing any homology to identified protein coding genes in other species (LOC100578137, LOC100577132, LOC100577221), and one protein coding gene showing homology with the probable *Apis cerana* serine/threonine-protein kinase dyrk2 gene (LOC413153). This was the gene containing 11 out of the 18 variants with SMA significance *p*≤0,001 in the locus of interest on chromosome 7. Of all protein kinases described in nature, most are classified as serine/threonine kinases, meaning that they catalyze the phosphorylation of serine or threonine residues on target proteins using ATP as a phosphate donor (92). Dual specificity tyrosine phosphorylation regulated kinase 2 (DYRK2) is categorized as a dual specificity kinase, indicating that it can act as both a tyrosine kinase and serine/threonine kinase (93, 94). DYRK2’s functions are well described in humans and other vertebrata, where it is involved in the regulation of the mitotic cell cycle, cellular growth and proliferation, proteostasis, p53-mediated apoptosis, and phosphorylation of histones (95–99). Direct physiological links between DYRK2 and viral infections in honey bees have not been reported yet. However, through injection of honey bee pupae with 10^4^ - 10^7^ DWV particles/bee, Mookhploy et al. (2020) showed that several genes involved in apoptosis were downregulated during the initial phase of viral infection, but upregulated in newly emerging DWV-infected adult bees (33). By consequence, if honey bee DYRK2 has a similar regulatory role in p53-mediated apoptosis as in humans, it may be involved in apoptosis-moderated reduction of DWV proliferation (33, 100–104). The current study reports the association of multiple variants in a single locus with the SOV trait in a single artificially created hybrid *Apis mellifera carnica* colony. Although the functions of some genes may be linked to viral infections or viral defense mechanisms in mammals, invertebrates or other insects, future functional or expression-related studies may elucidate the exact role of the discovered variants and genes in the establishment of the DWV-B viral status in our samples, or the SOV trait as a whole.

## Author contributions

**R.L.**, B.J.G.B., D.C.d.G., L.D.S., M.B., L.P. and A.G. conceived this research, designed the methodology, and interpreted the results. **R.L.**, M.B. and L.D.S. collected, screened, prepared and analyzed the samples. **R.L.** performed (statistical) data analysis and modeling. **R.L.** wrote the manuscript. **R.L.**, B.J.G.B., D.C.d.G., L.D.S., M.B., L.P. and A.G. reviewed the manuscript.

