## Supplementary Files for "A Putative Single-Locus Determinant of the Suppressed *In Ovo* Virus Infection (SOV) Trait in *Apis mellifera*"

**Supplementary materials**

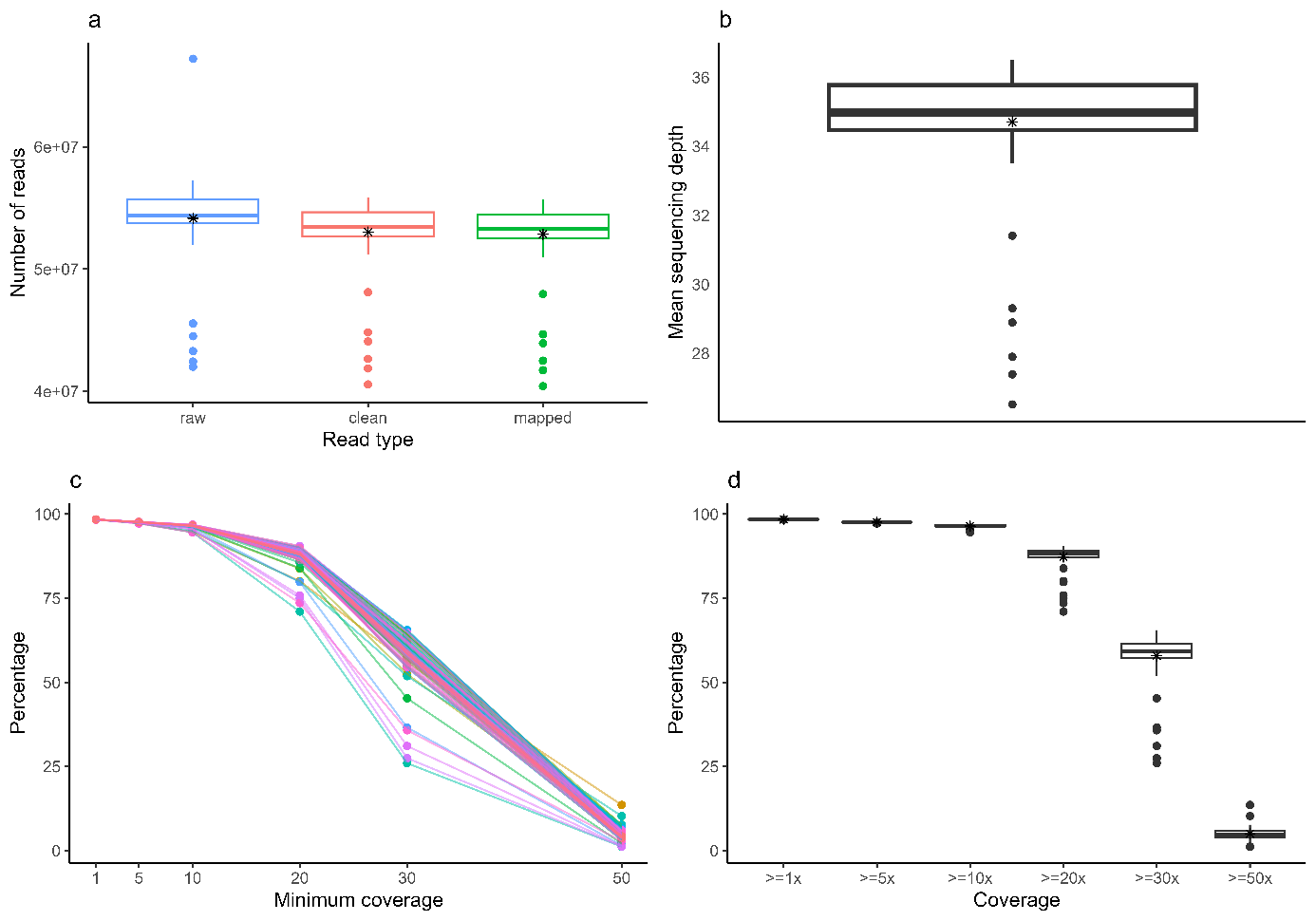

**Supplementary Figure S1. Overview of WGS- and mapping results. a.** Distribution of the number of raw reads, clean reads and mapped reads among the 88 samples. Averages are indicated with asterisks. **b.** Mean sequencing depths among the 88 samples. The average mean sequencing depth was 34,71x (asterisk). **c.** Relation between minimum coverage and percentage of target nucleotides for which the minimum coverage was reached. Dots are connected by sample. **d.** Idem, but distribution-wise. Plots are based on data from Supplementary Table S3.

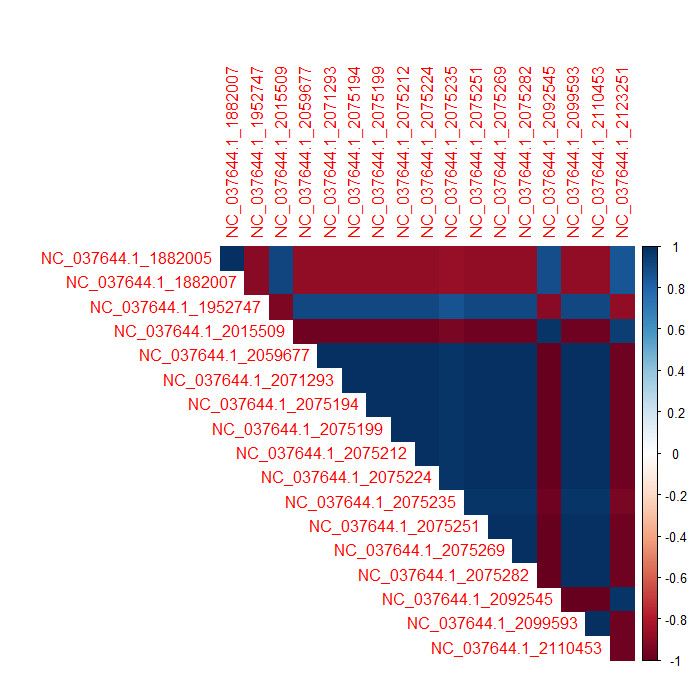

**Supplementary Figure S2. Allelic linkage disequilibria of the 7 indels and 11 SNPs located in the 241.246 nucleotide-wide locus on chromosome 7 (NC_037644.1).** Variants are coded as NC_037644.1_location. Blue (1): Wt-Wt and Vt-Vt linkage; red (-1): Wt-Vt linkage. Correlation plot is constructed for all samples with call for all 18 variants (N=67).

| Supplementary Table S1. Primer sets used for phenotyping by RT-qPCR. DWV: deformed wing virus; ABPV: acute bee paralysis virus; BQCV: black queen cell virus; SBV: sacbrood virus. DWV-FAM: primer set detecting the DWV complex. | | |
| --- | --- | --- |
| Target | **Forward primer** | **Reverse primer** |
| DWV-FAM (prescreening)  (105) | 5’-GGTAAGCGATGGTTGTTTG-3’ | 5’-CCGTGAATATAGTGTGAGG-3’ |
| DWV-B  (14) | 5’-TATCTTCATTAAAACCGCCAGGCT-3’ | 5’-CTTCCTCATTAACTGAGTTGTTGTC-3’ |
| β-ACTINE  (106) | 5’-AGGAATGGAAGCTTGCGGTA-3’ | 5’-AATTTTCATGGTGGATGGTGC-3’ |
| ABPV  (107) | 5’-TCATACCTGCCGATCAAG-3’ | 5’-CTGAATAATACTGTGCGTATC-3’ |
| BQCV  (108) | 5’-AGTGGCGGAGATGTATGC-3’ | 5’-GGAGGTGAAGTGGCTATATC-3’ |
| SBV  (108) | 5’-TTGGAACTACGCATTCTCTG-3’ | 5’-GCTCTAACCTCGCATCAAC-3’ |

| Supplementary Table S2. Overview of different filtering steps preceding the Single Marker Analyses (SMA). SNPs that met one of the following conditions were filtered by GATK VariantFiltration: QD < 2,0 \|\| FS > 60,0 \|\| MQ <40,0 \|\| MQRankSum < -12,5 \|\| ReadPosRankSum < -8,0. Indels that met one of the following conditions were filtered by GATK VariantFiltration: QD < 2,0 \|\| FS > 200,0 \|\| SOR > 10,0 \|\| MQRankSum < -12,5 \|\| ReadPosRankSum < -8,0. | | |
| --- | --- | --- |
| Filtering step | **Number of SNPs and indels remaining** | **Percentage of total (%)** |
| None (initial) | 2.203.547 | 100 |
| GATK VariantFiltration | 2.157.676 | 97,9 |
| Min. 2 alleles among the 88 samples | 1.301.515 | 59,1 |
| SNP or indel has call for at least 78 samples | **1.221.554** | 55,4 |

| **Supplementary Table S3. Overview of sequencing and mapping results.** Sample names are denoted as DWVB_N for negative samples, and DWVB_P for positive samples, followed by their sample number. | | | | | | | | | | | | |
| --- | --- | --- | --- | --- | --- | --- | --- | --- | --- | --- | --- | --- |
| **Sample** | **Clean reads** | **Mapping reads** | **Mapping rate** | **Properly paired reads** | **Properly paired ratio** | **Mean_depth** | **Coverage >= 1X** | **Coverage >= 5X** | **Coverage >= 10X** | **Coverage >= 20X** | **Coverage >= 30X** | **Coverage >= 50X** |
| DWVB_N10 | 53078668 | 52902895 | 99,7 | 52872328 | 99,6 | 34,7 | 98,4 | 97,7 | 96,7 | 88,7 | 59,6 | 4,2 |
| DWVB_N104 | 52700238 | 52521741 | 99,7 | 52487564 | 99,6 | 34,5 | 98,4 | 97,7 | 96,6 | 88,6 | 58,5 | 3,7 |
| DWVB_N106 | 55185134 | 55005606 | 99,7 | 54973972 | 99,6 | 36,1 | 98,5 | 97,7 | 96,8 | 90,5 | 63,8 | 4,8 |
| DWVB_N113 | 55781544 | 55593244 | 99,7 | 55557434 | 99,6 | 36,5 | 98,4 | 97,7 | 96,8 | 90,2 | 65 | 5,9 |
| DWVB_N118 | 53388676 | 53205097 | 99,7 | 53171206 | 99,6 | 34,9 | 98,4 | 97,7 | 96,7 | 89,3 | 61,1 | 4,4 |
| DWVB_N13 | 53031378 | 52852731 | 99,7 | 52820712 | 99,6 | 34,7 | 98,4 | 97,6 | 96,6 | 88,7 | 59,0 | 4,0 |
| DWVB_N149 | 53607754 | 53433324 | 99,7 | 53402794 | 99,6 | 35,1 | 98,4 | 97,6 | 96,6 | 89,1 | 60,2 | 4,1 |
| DWVB_N152 | 54890518 | 54715605 | 99,7 | 54685140 | 99,6 | 36,0 | 98,4 | 97,6 | 96,7 | 89,8 | 63,9 | 5,5 |
| DWVB_N156 | 55262082 | 55075007 | 99,7 | 55041368 | 99,6 | 36,2 | 98,4 | 97,6 | 96,4 | 89,3 | 65,4 | 6,6 |
| DWVB_N161 | 52411032 | 52244477 | 99,7 | 52216122 | 99,6 | 34,3 | 98,4 | 97,6 | 96,4 | 87,6 | 58,0 | 4,0 |
| DWVB_N163 | 54601600 | 54442291 | 99,7 | 54393266 | 99,6 | 35,7 | 98,4 | 97,3 | 94,7 | 80,1 | 55,9 | 13,6 |
| DWVB_N165 | 51160388 | 50993639 | 99,7 | 50963234 | 99,6 | 33,5 | 98,4 | 97,6 | 96,5 | 87,5 | 54,6 | 3,1 |
| DWVB_N166 | 53279784 | 53097871 | 99,7 | 53062346 | 99,6 | 34,9 | 98,4 | 97,6 | 96,6 | 88,8 | 61,1 | 4,9 |
| DWVB_N168 | 54552586 | 54374001 | 99,7 | 54342016 | 99,6 | 35,7 | 98,4 | 97,6 | 96,6 | 89,4 | 62,0 | 4,8 |
| DWVB_N171 | 52992594 | 52818327 | 99,7 | 52788418 | 99,6 | 34,7 | 98,4 | 97,6 | 96,6 | 89,2 | 60,9 | 4,4 |
| DWVB_N175 | 53308382 | 53133674 | 99,7 | 53102104 | 99,6 | 34,9 | 98,4 | 97,6 | 96,5 | 88,3 | 60,7 | 4,7 |
| DWVB_N180 | 53616610 | 53435603 | 99,7 | 53402622 | 99,6 | 35,1 | 98,4 | 97,6 | 96,5 | 88,6 | 60,7 | 4,9 |
| DWVB_N235 | 53252488 | 53070843 | 99,7 | 53041492 | 99,6 | 34,9 | 98,4 | 97,5 | 96,3 | 86,4 | 57,2 | 7,2 |
| DWVB_N240 | 51646196 | 51469577 | 99,7 | 51440714 | 99,6 | 33,9 | 98,4 | 97,5 | 96,0 | 83,8 | 52,5 | 6,1 |
| DWVB_N246 | 53125134 | 52949373 | 99,7 | 52922812 | 99,6 | 34,8 | 98,4 | 97,6 | 96,3 | 86,8 | 57,3 | 6,1 |
| DWVB_N251 | 55418576 | 55239646 | 99,7 | 55213412 | 99,6 | 36,3 | 98,4 | 97,6 | 96,4 | 88,4 | 61,4 | 7,7 |
| DWVB_N257 | 55090526 | 54904129 | 99,7 | 54870166 | 99,6 | 36,1 | 98,4 | 97,6 | 96,6 | 89,3 | 63,2 | 5,8 |
| DWVB_N265 | 55705742 | 55541582 | 99,7 | 55517778 | 99,7 | 36,4 | 98,4 | 97,6 | 96,3 | 87,9 | 64,2 | 7,6 |
| DWVB_N274 | 54739388 | 54572711 | 99,7 | 54546642 | 99,7 | 35,8 | 98,4 | 97,6 | 96,6 | 88,5 | 61,6 | 5,8 |
| DWVB_N287 | 51776944 | 51611541 | 99,7 | 51589034 | 99,6 | 33,9 | 98,4 | 97,6 | 96,5 | 87,8 | 56,6 | 3,7 |
| DWVB_N288 | 55839582 | 55667150 | 99,7 | 55641430 | 99,7 | 36,5 | 98,4 | 97,7 | 96,7 | 90,0 | 64,4 | 5,7 |
| DWVB_N291 | 55442418 | 55277709 | 99,7 | 55253634 | 99,7 | 36,3 | 98,4 | 97,6 | 96,6 | 89,6 | 64,5 | 6,4 |
| DWVB_N292 | 54173480 | 54010980 | 99,7 | 53987162 | 99,7 | 35,4 | 98,4 | 97,6 | 96,4 | 87,4 | 59,8 | 5,6 |
| DWVB_N297 | 52566780 | 52409333 | 99,7 | 52384054 | 99,7 | 34,4 | 98,4 | 97,6 | 96,5 | 87,3 | 57,2 | 4,5 |
| DWVB_N3 | 53891924 | 53712217 | 99,7 | 53679000 | 99,6 | 35,3 | 98,4 | 97,6 | 96,4 | 87,8 | 60,6 | 5,0 |
| DWVB_N302 | 48106422 | 47954397 | 99,7 | 47932124 | 99,6 | 31,4 | 98,4 | 97,5 | 96,1 | 84,0 | 45,3 | 2,1 |
| DWVB_N303 | 55064204 | 54881746 | 99,7 | 54855798 | 99,6 | 36,0 | 98,4 | 97,7 | 96,7 | 89,3 | 61,8 | 5,1 |
| DWVB_N305 | 54138892 | 53979481 | 99,7 | 53948808 | 99,7 | 35,4 | 98,4 | 97,6 | 96,4 | 86,6 | 58,8 | 6,7 |
| DWVB_N314 | 54855554 | 54701514 | 99,7 | 54672076 | 99,7 | 35,9 | 98,4 | 97,6 | 96,5 | 87,2 | 59,7 | 6,8 |
| DWVB_N316 | 53630042 | 53485367 | 99,7 | 53459356 | 99,7 | 35,1 | 98,4 | 97,6 | 96,2 | 85,6 | 57,3 | 6,9 |
| DWVB_N37 | 52728146 | 52549335 | 99,7 | 52516720 | 99,6 | 34,5 | 98,4 | 97,6 | 96,5 | 87,6 | 58,6 | 4,4 |
| DWVB_N55 | 55095796 | 54920507 | 99,7 | 54894520 | 99,6 | 36,2 | 98,4 | 97,6 | 96,4 | 88,0 | 60,4 | 7,5 |
| DWVB_N60 | 54141426 | 53959185 | 99,7 | 53929584 | 99,6 | 35,5 | 98,4 | 97,5 | 96,4 | 88,1 | 58,9 | 6,5 |
| DWVB_N62 | 53833092 | 53661423 | 99,7 | 53638198 | 99,6 | 35,3 | 98,4 | 97,5 | 96,1 | 86,8 | 59,5 | 7,3 |
| DWVB_N69 | 55152342 | 54974233 | 99,7 | 54942548 | 99,6 | 36,1 | 98,4 | 97,7 | 96,7 | 89,3 | 62,7 | 5,4 |
| DWVB_N82 | 40566642 | 40420587 | 99,7 | 40391432 | 99,6 | 26,5 | 98,4 | 97,3 | 94,6 | 71,0 | 26,0 | 1,2 |
| DWVB_N84 | 52243854 | 52096636 | 99,7 | 52061074 | 99,7 | 34,3 | 98,3 | 97,3 | 95,0 | 79,9 | 51,9 | 10,3 |
| DWVB_N96 | 53685018 | 53502359 | 99,7 | 53468806 | 99,6 | 35,1 | 98,4 | 97,7 | 96,7 | 89,1 | 60,5 | 4,5 |
| DWVB_N97 | 55411236 | 55219257 | 99,7 | 55180168 | 99,6 | 36,3 | 98,4 | 97,7 | 96,7 | 89,9 | 63,9 | 5,7 |
| DWVB_P101 | 53555984 | 53379396 | 99,7 | 53348138 | 99,6 | 35,1 | 98,4 | 97,6 | 96,3 | 86,5 | 57,5 | 7,7 |
| DWVB_P105 | 52244698 | 52067352 | 99,7 | 52032102 | 99,6 | 34,2 | 98,4 | 97,6 | 96,5 | 87,0 | 55,0 | 3,6 |
| DWVB_P111 | 53834688 | 53659640 | 99,7 | 53626718 | 99,6 | 35,2 | 98,4 | 97,6 | 96,7 | 89,2 | 61,0 | 4,7 |
| DWVB_P112 | 52144056 | 51974021 | 99,7 | 51942878 | 99,6 | 34,1 | 98,4 | 97,6 | 96,5 | 87,7 | 57,7 | 4,1 |
| DWVB_P129 | 54699882 | 54522543 | 99,7 | 54488074 | 99,6 | 35,8 | 98,4 | 97,6 | 96,5 | 89,0 | 61,8 | 5,1 |
| DWVB_P147 | 55744570 | 55558175 | 99,7 | 55523474 | 99,6 | 36,5 | 98,4 | 97,7 | 96,6 | 89,8 | 65,6 | 6,3 |
| DWVB_P158 | 53262634 | 53084933 | 99,7 | 53051112 | 99,6 | 34,8 | 98,4 | 97,6 | 96,6 | 89,1 | 59,9 | 4,2 |
| DWVB_P172 | 53541398 | 53360390 | 99,7 | 53324574 | 99,6 | 35,0 | 98,4 | 97,6 | 96,5 | 88,5 | 60,3 | 4,6 |
| DWVB_P178 | 52918814 | 52740235 | 99,7 | 52704778 | 99,6 | 34,6 | 98,4 | 97,6 | 96,6 | 88,5 | 57,1 | 3,5 |
| DWVB_P182 | 53099276 | 52916152 | 99,7 | 52873244 | 99,6 | 34,7 | 98,4 | 97,6 | 96,5 | 87,6 | 58,3 | 4,8 |
| DWVB_P183 | 54184820 | 54011019 | 99,7 | 53979384 | 99,6 | 35,5 | 98,4 | 97,6 | 96,7 | 89,6 | 62,4 | 5,3 |
| DWVB_P184 | 51388626 | 51216523 | 99,7 | 51182368 | 99,6 | 33,6 | 98,4 | 97,6 | 96,6 | 87,7 | 54,3 | 3,2 |
| DWVB_P185 | 55748246 | 55552762 | 99,7 | 55508696 | 99,6 | 36,4 | 98,4 | 97,7 | 96,7 | 90,1 | 65,2 | 6,5 |
| DWVB_P188 | 54050306 | 53886865 | 99,7 | 53862174 | 99,7 | 35,5 | 98,4 | 97,5 | 96,3 | 87,0 | 58,0 | 6,5 |
| DWVB_P189 | 44822724 | 44666843 | 99,7 | 44636364 | 99,6 | 29,3 | 98,4 | 97,5 | 95,9 | 79,8 | 36,6 | 1,6 |
| DWVB_P190 | 52399620 | 52212085 | 99,7 | 52174432 | 99,6 | 34,3 | 98,4 | 97,7 | 96,7 | 88,2 | 57,8 | 4,0 |
| DWVB_P191 | 53004256 | 52832304 | 99,7 | 52801770 | 99,6 | 34,7 | 98,4 | 97,6 | 96,7 | 88,8 | 58,6 | 3,9 |
| DWVB_P192 | 54597400 | 54412580 | 99,7 | 54379128 | 99,6 | 35,7 | 98,4 | 97,7 | 96,7 | 89,2 | 61,4 | 4,7 |
| DWVB_P193 | 53062348 | 52880958 | 99,7 | 52846896 | 99,6 | 34,7 | 98,4 | 97,6 | 96,7 | 88,7 | 58,2 | 3,7 |
| DWVB_P194 | 55215602 | 55021350 | 99,7 | 54983622 | 99,6 | 36,1 | 98,4 | 97,7 | 96,8 | 89,6 | 63,6 | 5,8 |
| DWVB_P195 | 53340046 | 53159007 | 99,7 | 53120134 | 99,6 | 34,9 | 98,4 | 97,7 | 96,6 | 87,5 | 59,5 | 5,8 |
| DWVB_P196 | 52976470 | 52797505 | 99,7 | 52764088 | 99,6 | 34,7 | 98,4 | 97,7 | 96,7 | 89,1 | 59,0 | 3,9 |
| DWVB_P199 | 53999742 | 53807602 | 99,6 | 53768880 | 99,6 | 35,3 | 98,4 | 97,7 | 96,7 | 89,5 | 60,1 | 4,0 |
| DWVB_P200 | 41872136 | 41726042 | 99,7 | 41697202 | 99,6 | 27,4 | 98,3 | 97,4 | 95,6 | 75,0 | 27,5 | 1,2 |
| DWVB_P202 | 55544382 | 55358179 | 99,7 | 55321956 | 99,6 | 36,4 | 98,4 | 97,7 | 96,9 | 90,5 | 64,8 | 5,8 |
| DWVB_P204 | 42652816 | 42503491 | 99,7 | 42475348 | 99,6 | 27,9 | 98,4 | 97,4 | 95,5 | 75,9 | 31,1 | 1,4 |
| DWVB_P205 | 51744032 | 51565509 | 99,7 | 51531294 | 99,6 | 33,9 | 98,4 | 97,6 | 96,5 | 86,8 | 56,1 | 4,0 |
| DWVB_P206 | 55216578 | 55035066 | 99,7 | 55000286 | 99,6 | 36,1 | 98,4 | 97,7 | 96,7 | 88,9 | 61,7 | 5,8 |
| DWVB_P207 | 55316558 | 55136942 | 99,7 | 55101830 | 99,6 | 36,2 | 98,4 | 97,6 | 96,6 | 88,7 | 62,0 | 6,0 |
| DWVB_P21 | 53627968 | 53446316 | 99,7 | 53408670 | 99,6 | 35,1 | 98,4 | 97,6 | 96,6 | 88,5 | 59,3 | 4,6 |
| DWVB_P214 | 51969618 | 51797984 | 99,7 | 51773040 | 99,6 | 34,1 | 98,4 | 97,5 | 96,3 | 86,4 | 55,4 | 5,0 |
| DWVB_P22 | 51783404 | 51608782 | 99,7 | 51574082 | 99,6 | 33,9 | 98,4 | 97,6 | 96,5 | 86,9 | 54,5 | 3,4 |
| DWVB_P220 | 44085768 | 43929007 | 99,6 | 43902512 | 99,6 | 28,9 | 98,3 | 97,2 | 94,6 | 73,6 | 35,8 | 2,8 |
| DWVB_P230 | 53788446 | 53612648 | 99,7 | 53587118 | 99,6 | 35,3 | 98,4 | 97,6 | 96,5 | 88,8 | 59,4 | 5,4 |
| DWVB_P24 | 54633936 | 54449552 | 99,7 | 54412728 | 99,6 | 35,8 | 98,4 | 97,7 | 96,8 | 89,9 | 62,1 | 5,0 |
| DWVB_P25 | 52994000 | 52813754 | 99,7 | 52777462 | 99,6 | 34,7 | 98,4 | 97,7 | 96,6 | 88,0 | 58,1 | 4,4 |
| DWVB_P26 | 51788278 | 51611400 | 99,7 | 51576542 | 99,6 | 33,9 | 98,4 | 97,6 | 96,5 | 86,7 | 55,2 | 3,9 |
| DWVB_P28 | 52526738 | 52340164 | 99,6 | 52305840 | 99,6 | 34,4 | 98,4 | 97,7 | 96,5 | 87,9 | 58,6 | 4,1 |
| DWVB_P30 | 53165966 | 52975653 | 99,6 | 52937206 | 99,6 | 34,8 | 98,4 | 97,7 | 96,7 | 88,8 | 58,4 | 3,9 |
| DWVB_P40 | 53638844 | 53448412 | 99,6 | 53409214 | 99,6 | 35,2 | 98,4 | 97,6 | 96,6 | 88,7 | 58,8 | 4,1 |
| DWVB_P41 | 53130250 | 52945939 | 99,7 | 52908340 | 99,6 | 34,8 | 98,4 | 97,6 | 96,6 | 88,3 | 57,4 | 3,8 |
| DWVB_P43 | 52516472 | 52345338 | 99,7 | 52314252 | 99,6 | 34,4 | 98,4 | 97,7 | 96,6 | 87,9 | 57,1 | 3,8 |
| DWVB_P46 | 53110758 | 52931936 | 99,7 | 52900104 | 99,6 | 34,8 | 98,4 | 97,6 | 96,5 | 87,9 | 58,5 | 3,8 |
| DWVB_P47 | 52760422 | 52579565 | 99,7 | 52544218 | 99,6 | 34,6 | 98,4 | 97,7 | 96,5 | 88,2 | 59,4 | 4,3 |
| **Average** | **53099907** | **52846249** | **99,7** | **52814130** | **99,6** | **34,7** | **98,4** | **97,6** | **96,4** | **87,3** | **57,9** | **5,0** |

| Supplementary Table S4. Overview of the most significant SNP and indel per chromosome in the SMA (N = 1.221.554). Locations and gene names refer to the reference genome Amel_HAv3.1. Uncorrected p-values are reported. Undescribed genes indicate that the variant was not located in a gene. | | | | |
| --- | --- | --- | --- | --- |
| Chromosome | **Variant** | **Location** | **Gene** | **Significance** |
| 1 | SNP | 13.068.823 |  | 0,00248 |
|  | Indel | 13.917.351 |  | 0,000771 |
| 2 | SNP | 9.606.004 | LOC413350 - lachesin | 0,00296 |
|  | Indel | 10.883.424 | LOC100577670 - uncharacterized | 0,000728 |
| 3 | SNP | 13.014.519 | LOC413503 - TWiK family of potassium channels protein 18-like | 0,00597 |
|  | Indel | 922.528 | LOC100577185 - uncharacterized | 0,000150 |
| 4 | SNP | 10.975.793 | LOC411018 - kinesin 11 | 0,00797 |
|  | Indel | 12.629.047 | LOC408775 - synaptotagmin binding cytoplasmic RNA interacting protein | 0,000201 |
| 5 | SNP | 9.992.227 |  | 0,00174 |
|  | Indel | 10.632.879 | LOC100578865 - CABIT domain-containing protein serrano | 0,00100 |
| 6 | SNP | 108.502 | LOC100578680 - bloated | 0,00227 |
|  | Indel | 6.778.691 | LOC412917 - probable serine/threonine-protein kinase dyrk2 | 0,000898 |
| 7 | SNP | 2.059.677 | LOC413153 – uncharacterized | 0,000419 |
|  | Indel | 1.952.747 |  | 0,000108 |
| 8 | SNP | 11.459.866 | LOC551016 - mRNA-midasin | 0,000659 |
|  | Indel | 9.616.930 |  | 0,000659 |
| 9 | SNP | 11.404.110 | LOC409409 - serine/threonine-protein kinase dyf-5 | 0,00212 |
|  | Indel | 11.404.115 |  | 0,000668 |
| 10 | SNP | 602.271 | LOC413242 - N-deacetylase and N-sulfotransferase sfl | 0,00387 |
|  | Indel | 10.219.150 | LOC409315 - protein interacting with PRKCA 1 | 0,00103 |
| 11 | SNP | 1.397.351 | LOC408339 – uncharacterized  LOC102654166 - trichohyalin | 0,00344 |
|  | Indel | 6.706.915 | LOC724942 - putative phosphatidate phosphatase | 0,000210 |
| 12 | SNP | 4.988.673 |  | 0,00458 |
|  | Indel | 8.161.396 | LOC410166 - myelin regulatory factor-like protein | 0,000548 |
| 13 | SNP | 9.678.320 |  | 0,00434 |
|  | Indel | 5.349.407 | forkhead box P | 0,000791 |
| 14 | SNP | 6.252.297 | LOC726153 - uncharacterized | 0,000718 |
|  | Indel | 2.886.600 | LOC413213 - tubulin polyglutamylase TTLL4 | 0,00313 |
| 15 | SNP | 6.608.885 | LOC552844 - synaptic transmission protein complexin | 0,00349 |
|  | Indel | 53.099 |  | 0,00375 |
| 16 | SNP | 97.158 | LOC107965756 - uncharacterized | 0,00330 |
|  | Indel | 5.337.229 | LOC550708 - mind bomb 1  LOC107965683 - uncharacterized | 0,00961 |

| Supplementary Table S5. Overview of the 18 variants with (non- Bonferroni-corrected) significance *p*≤0,001 in the 241.246 nucleotide-wide locus on chromosome 7. Positions, gene codes and gene names refer to the reference genome Amel_HAv3.1. 3’ UTR: three prime untranslated region. | | | | | | |
| --- | --- | --- | --- | --- | --- | --- |
| Position | **Variant** | **WT** | **VT** | **Intron/exon/3’ UTR** | **Gene code** | **Gene** |
| 1.882.005 | Indel | A | AAGAATTTAAAAATACGAGAAT | Intron | LOC411586 | GAS2-like protein pickled eggs |
| 1.882.007 | Indel | G | GGAATACGAATTTAAGA |  |  |  |
| 1.952.747 | Indel | AATAT | A, AAT | - | - | - |
| 2.015.509 | SNP | G | A | Intron | LOC413153 | Uncharacterized |
|  |  |  |  |  | LOC100577098 | Enolase-phosphatase E1 |
| 2.059.677 | SNP | C | T | Intron | LOC413153 | Uncharacterized |
| 2.071.293 | Indel | AC | A | Intron |  |  |
| 2.075.194 | SNP | A | G | Intron |  |  |
| 2.075.199 | SNP | T | C |  |  |  |
| 2.075.212 | SNP | G | A |  |  |  |
| 2.075.224 | SNP | T | C |  |  |  |
| 2.075.235 | Indel | A | AATATATATATATATAT, AACATATATATATATAT |  |  |  |
| 2.075.251 | SNP | C | T, A |  |  |  |
| 2.075.269 | SNP | T | C |  |  |  |
| 2.075.282 | SNP | T | C |  |  |  |
| 2.092.545 | SNP | A | G | Intron | LOC100577026 | mitochondrial amidoxime-reducing component 1-like |
| 2.099.593 | Indel | A | AT | Intron | LOC551405 | 4-nitrophenyl-phosphatase |
| 2.110.453 | Indel | C | CTT, CT | 3’ UTR | LOC411748 | protein kinase C-binding protein NELL1 |
| 2.123.251 | SNP | G | A | Intron | LOC411748 | protein kinase C-binding protein NELL1 |

| Supplementary Table S6. Overview of elastic net regression results. During elastic net regression analyses, unaltered variables were all SNPs and indels with a call for all 88 samples (N = 1.149.592) and the phenotype of the samples (DWV-B negative (0); DWV-B positive (1)). For optimal lambda and alpha selection, alpha was altered from 1 to 0,1 in steps of 0,1 and lambda was tested repetitively for 100 different values. For each value of alpha, the number of included variants, the Log_10_ of the number of variants, and percentages of correct prediction by the models at λ_min_ and λ_1.se_ are represented. | | | | |
| --- | --- | --- | --- | --- |
| Alpha | **Lambda** | **N variants** | **Log_10_ (N variants)** | **Correct prediction (%)** |
| 1  (LASSO only) | 1se | 1 | 0 | 68,2 |
|  | min | 1 | 0 | 68,2 |
| 0,9 | 1se | 1 | 0 | 68,2 |
|  | min | 1 | 0 | 68,2 |
| 0,8 | 1se | 1 | 0 | 68,2 |
|  | min | 1 | 0 | 68,2 |
| 0,7 | 1se | 1 | 0 | 68,2 |
|  | min | 1 | 0 | 68,2 |
| 0,6 | 1se | 5 | 0,698970004 | 69,3 |
|  | min | 7 | 0,84509804 | 71,6 |
| 0,5 | 1se | 6 | 0,77815125 | 68,2 |
|  | min | 10 | 1 | 71,6 |
| 0,4 | 1se | 6 | 0,77815125 | 69,3 |
|  | min | 10 | 1 | 71,6 |
| 0,3 | 1se | 6 | 0,77815125 | 69,3 |
|  | min | 11 | 1,041392685 | 71,6 |
| 0,2 | 1se | 7 | 0,84509804 | 69,3 |
|  | min | 7 | 0,84509804 | 69,3 |
| 0,1 | 1se | 10 | 1 | 69,3 |
|  | min | 526 | 2,720985744 | 70,5 |

| Supplementary Table S7. Indel (NC_037644.1_1952747) -associated genotypes and alleles of the remaining 17 SMA-significant variants located along the 241.246 nucleotide-wide locus on chromosome 7. The ‘associated genotype’ column represents the genotype of the represented genetic variant that is associated with the variant type alleles of the indel NC_037644.1_1952747. N represents the number of samples included in the Fisher’s Exact Tests (excluding samples with ‘NA’ genotypes from variant calling). | | | | | |
| --- | --- | --- | --- | --- | --- |
| Variant | **Location** | **Associated genotype** | **Associated allele** | **Gene** | **N** |
| Indel | 1.882.005 | Wt | A | GAS2-like protein pickled eggs (LOC411586) | 88 |
| Indel | 1.882.007 | Wt | G |  | 88 |
| SNP | 2.015.509 | Wt | G | Uncharacterized (LOC413153)  Enolase-phosphatase E1 (LOC100577098) | 83 |
| SNP | 2.059.677 | Vt | T | Uncharacterized (LOC413153) | 85 |
| Indel | 2.071.293 | Vt | A |  | 87 |
| SNP | 2.075.194 | Vt | G |  | 87 |
| SNP | 2.075.199 | Vt | C |  | 87 |
| SNP | 2.075.212 | Vt | A |  | 87 |
| SNP | 2.075.224 | Vt | C |  | 87 |
| Indel | 2.075.235 | Vt | AATATATATATATATAT, AACATATATATATATAT |  | 85 |
| SNP | 2.075.251 | Vt | T, A |  | 82 |
| SNP | 2.075.269 | Vt | C |  | 80 |
| SNP | 2.075.282 | Vt | C |  | 80 |
| SNP | 2.092.545 | Wt | A | mitochondrial amidoxime-reducing component 1-like (LOC100577026) | 87 |
| Indel | 2.099.593 | Vt | AT | 4-nitrophenylphosphatase (LOC551405) | 87 |
| Indel | 2.110.453 | Vt | CTT, CT | protein kinase C-binding protein NELL1 (LOC411748) | 88 |
| SNP | 2.123.251 | Wt | G | protein kinase C-binding protein NELL1 (LOC411748) | 85 |
